# Gametophytes and embryo ontogeny: understanding the reproductive calendar of *Cypripedium japonicum* Thunb. (Cypripedoideae, Orchidaceae)

**DOI:** 10.1101/738799

**Authors:** Balkrishna Ghimire, Sung Won Son, Jae Hyun Kim, Mi-Jin Jeong

## Abstract

Among the flowering plants, the gametophyte development and reproductive biology of orchids is particularly poorly understood. *Cypripedium japonicum* is a perennial herb, native to East Asia. Due to its limited distribution, the species is included in the Endangered category of the IUCN Red List. Light microscopy and SEM methods were used to study the development of the gametes and embryo. The complete reproductive cycle was developed based on our observations. Anther development begins under the soil and meiosis of pollen cells begins 3 weeks before anthesis, possibly during early April. The megaspore mother cells develop just after pollination in early May and mature in mid–late June. The pattern of embryo sac formation is bisporic and there are six nuclei. Triple fertilization results in the endosperm nucleus. A globular embryo is formed after multiple cell division and 9 weeks after pollination the entire embryo sac is occupied by embryo. Overall comparisons of the features of gametophyte and embryo development in *C. japonicum* suggest that previous reports on the embryology of *Cypripedium* are not sufficient to characterize the entire genus. Based on the available information a reproductive calendar showing the key reproductive events leading to embryo formation has been prepared.

**Highlight:** Manual pollination, reproductive biology and seed development process in *Cypripedium japonicum* Thunb., a lady’s slipper orchid endemic to East Asia

## Introduction

The family Orchidaceae is one of the largest families of flowering plants, along with the Asteraceae (Dressler, 1993; Mabberley, 2008). The orchid taxa exhibit an unusual pattern of embryo development and the diverse morphology of the suspensor is one of the most distinctive features in the whole family (Arditti, 1992; Dressler, 1993; Lee *et al*., 2006). The characteristic suspensor development, patterns of polyembryony, and vast diversity of megagametophyte development in the orchids provide continual fascination for plant biologists investigating the embryology of this family (Swamy, 1949a, 1949b; Sood, 1985; Young *et al*., 1996; Lee *et al*., 2006). The extraordinarily diverse pattern of suspensor morphology in orchids motivated Swamy (1949b) to conceive a classification scheme for embryo development in the Orchidaceae. In addition, the anthers usually develop much earlier than the ovule in orchid species; at pollination when anthers shed mature pollen grains containing a fully developed male gamete the ovule has yet to produce megaspore mother cells (MMCs; (Swamy, 1943; Wirth and Withner, 1959). It has been suggested that ovule development is activated by pollination, after which it takes several weeks to produce a female gamete (Arditti, 1992; Yeung and Law, 1997). Zhang and O’Neill (1993) believed that the widespread abundance of prolonged period of ovule development in the family make the Orchidaceae an intriguing prospect for embryological study.

The genus *Cypripedium* L. is one of the five genera of the subfamily Cypripedioideae, commonly known as the lady’s slippers. The members of this group are mostly distributed in temperate and certain tropical zones of the Northern Hemisphere (Dressler, 1993; Cribb, 1997). The beautiful flowers and hardiness of these plants make them a very popular garden plant with high ornamental and commercial value and thus are attractive to botanists and floriculturists alike (Dressler, 1993; Jong, 2002). All the species of this group have a pouch-like lip, two fertile stamens, a shield-like staminoide, and a synsepal with fused lateral sepals, and this specific flower morphology does not offer an easy reward to pollinators (Lindely, 1840; Pridgeon *et al*. 1999). A unique feature of Cypripedioideae orchids is the one-way trap flower which offers a fixed and one-way route for pollinators (see Li *et al*., 2008). In spite of this fixed pollination route, slipper orchids demonstrate a remarkable degree of diversity in their pollinator and pollination mechanisms (Stoutamire, 1967; Nilsson, 1979; Sugiura *et al*., 2001; Banziger and Sun, 2005; Li *et al*., 2008). We performed manual pollination in our research; pollinators and pollination mechanisms are thus beyond the scope of this study.

The genus *Cypripedium* comprises about 45–50 species which are mostly found in temperate zones of the Northern Hemisphere, mainly in temperate Asia and North America, extending to the Himalayan regions and Central America (Cribb, 1997; Wu *et al*., 2009; Guo *et al*., 2012; WCSP, 2019). The center of diversity of the genus lies in Eastern Asia where 38 species have been reported; in particular, China alone has an astonishing diversity of the genus, with 36 species of which 25 species are endemic (Jong, 2002; Wu *et al*., 2009). Many species of *Cypripedium* in North America and Europe have become rare due to over-exploitation and illegal trade in past centuries (Case *et al*., 1998; Ramsay and Stewart, 1998). The condition for *Cypripedium* in northeastern Asia is no different. Some 24 Asian *Cypripedium* species are currently included in the Endangered or Critically Endangered categories of the IUCN Red List (IUCN, 2019). In addition, the Convention on International Trade in Endangered Species of Wild Fauna and Flora (CITES) currently lists all *Cypripedium* species in CITES Appendix II (CITES, 2007).

*C. japonicum* Thunb. is one species of endangered East Asian endemic lady’s slipper orchid, which in particular is distributed in China, Japan, and Korea. It grows in damp and humus-rich soil in forest or on shady slopes along ravines at an altitude of 1000–2000 m (Chen *et al*., 1999). In Korea, this species is extremely rare, and fewer than ten populations are recognized from four isolated localities in old and wet deciduous forest on mountain hillsides with an elevation of approximately 500–700 m (Chung *et al*., 2009). Due to its limited distribution with small populations in natural habitat this species has been categorized as Critically Endangered (although it is Endangered globally) at the national level (Korea National Arboretum, 2008). The South Korean government has designated the species as ‘Threatened to Extinct: the first grade (I) for preservation’ and it is protected under the Wildlife Protection Act of Korea (Ministry of Environment in Korea, 2012). Correspondingly, in China and Japan the species is at a high risk of extinction due to habitat loss and anthropogenic activities like over-collection for horticultural and medicinal purposes (Ministry of the Environment in Japan, 2015; Qian *et al*., 2014).

An empirical knowledge of plant reproductive biology, particularly of rare orchids, could help us determine whether inadequacy in the recruitment cycle is controlling successful reproduction and constraining population growth (Gale, 2007). A few recent efforts have been made to study the pollination biology and reproductive characteristics of *C. japonica* (Suetsugu and Fukushima, 2013; Sun *et al*., 2009). In the early 20th century Pace (1907) made a comprehensive account of fertilization in the genus *Cypripedium* although *C. japonica* was not examined. Later, Sood and Rao (1988) studied embryology of *C. cordigerum* and confirmed that the embryological characters and pericarp structure in this species were typical orchidaceous types. Although the majority of orchids follow a monosporic pattern of embryo sac development (Vij and Sharma, 1986; Johri *et al*., 1992), bisporic pattern has been described for *Cypripedium* and *Paphiopedilum* (Duncan and Curtis, 1942; Carlson, 1954). While there is no structural or developmental examination of antipodal cells in Pace’s (1907) report, Poddubnaya-Arnoldi (1967) described the well-developed antipodal cells which undergo secondary multiplication in some *Cypripedium* species. In the past few years, several studies have been carried out focusing on the embryology of different orchid species (Lee *et al*., 2006, 2008; Lee and Young, 2012; Wang *et al*., 2016; Li *et al*., 2016). However, little is known about the embryology and reproduction biology of *C. japonicum*. Knowledge of gametophyte, embryo, and seed development is essential for the understanding of successful reproduction in this endangered orchid, and for formulating a restoration strategy.

We investigated the detailed embryology and seed formation processes in this popular lady’s slipper orchid. The primary objectives of this study were: (1) to investigate male and female gametophyte development in artificially pollinated *C. japonicum*, (2) to comprehend the fertilization, embryo, and seed development processes in *C. japonicum*, and (3) to develop a complete reproductive calendar of the species showing key reproductive events leading to seed development.

## Materials and methods

### Plant materials

Several clumps of *Cypripedium japonicum* were grown inside the fence near one of the six natural populations in the Korea (see Chung *et al*., 2009). To ensure quality fruit set, flowers were manually cross-pollinated by transferring pollen of one flower onto the stigma of another flower, with an effort not to limit cross-pollination to the same clump. A whole anther was detached from a flower with a pointed pin set, rinsed with 70% ethanol prior to use, and attached to the stigma of another flower. To avoid unwanted pollination by insects the pollinated flowers were covered by a finely netted nylon bag. For a comprehensive study of embryology, developing flowers and fruits were collected at regular time intervals before and after the pollination. Sample collections were started at the end of March, when new shoots start to arise, until early September, when fruit is completely ripened and start to rupture its wall for seed dispersal. In early collections, particularly during March, which were collected mainly to study male gametophyte development, all the developing shoots were underneath the soil surface. All the collected samples were fixed in formalin/acetic acid/50% ethyl alcohol (FAA) in the ratio 90:5:5 per 100 ml for at least 3–5 days and preserved in 50% ethyl alcohol.

### Light microscopy

Preserved flower buds, open flowers and fruits were dehydrated in an ethyl alcohol series (50, 70, 80, 90, 95, and 100%) for at least 24 h in each stage. After complete dehydration, the samples were passed through an infiltration solution of alcohol/Technovit combinations (3:1, 1:1, 1:3, and 100% Technovit) and then embedded in polymerization solution prepared by mixing Technovit 7100 resin stock solution with Technovit 7100 harder II. Histological blocks were prepared from each embedded material with Histoblock and Technovit 3040 so that the mold could be removed and the material sectioned. Serial sections of 4–6 µm thickness were cut using a Leica RM2255 rotary microtome (Leica Microsystems GmbH, Germany) with disposable blades, then stuck onto histological slides and dried using an electric slide warmer for 12 h. Dried slides were stained with 0.1% Toluidine blue O for 60–90 s, rinsed with running water, and again dried with an electric slide warmer for more than 6 h to remove water. The stained slides were then mounted with Entellan (Merck Co., Germany) and compressed using metal blocks for 2 days to remove air bubbles. These permanent slides were observed under an AXIO Imager A1 light microscope (Carl Zeiss, Germany). Photomicrographs were taken with an attached camera system.

### Scanning electron microscopy

For scanning electron microscopy, young and mature pollinia and ovaries were dehydrated by ethanol series, dried using a Samdri – PVT – 3D critical point dryer (Tousimis Co., USA) and sputtered with gold coating in a KIC-IA COXEM Ion-Coater (COXEM Co., Korea). Scanning electron microscopic imaging was carried out with a COXEM CX-100S scanning electron microscope at 20 kV in the seed testing laboratory of the Korea National Arboretum.

## Results

Several new buds developed from the rhizome (Fig. 1A). A single flower is borne at the tip of the initial shoot. Flower development starts when the whole plant is still under the soil surface. The mature flower is large and showy with greenish yellow petals with purple spots at the base. The lip is obovoid or ellipsoidal, pouch-like, and yellowish with purple spots (Fig. 1B). Flowers were hand-pollinated and young to mature fruits were collected for the experiment (Fig. 1C, D).

**Fig. 1.**
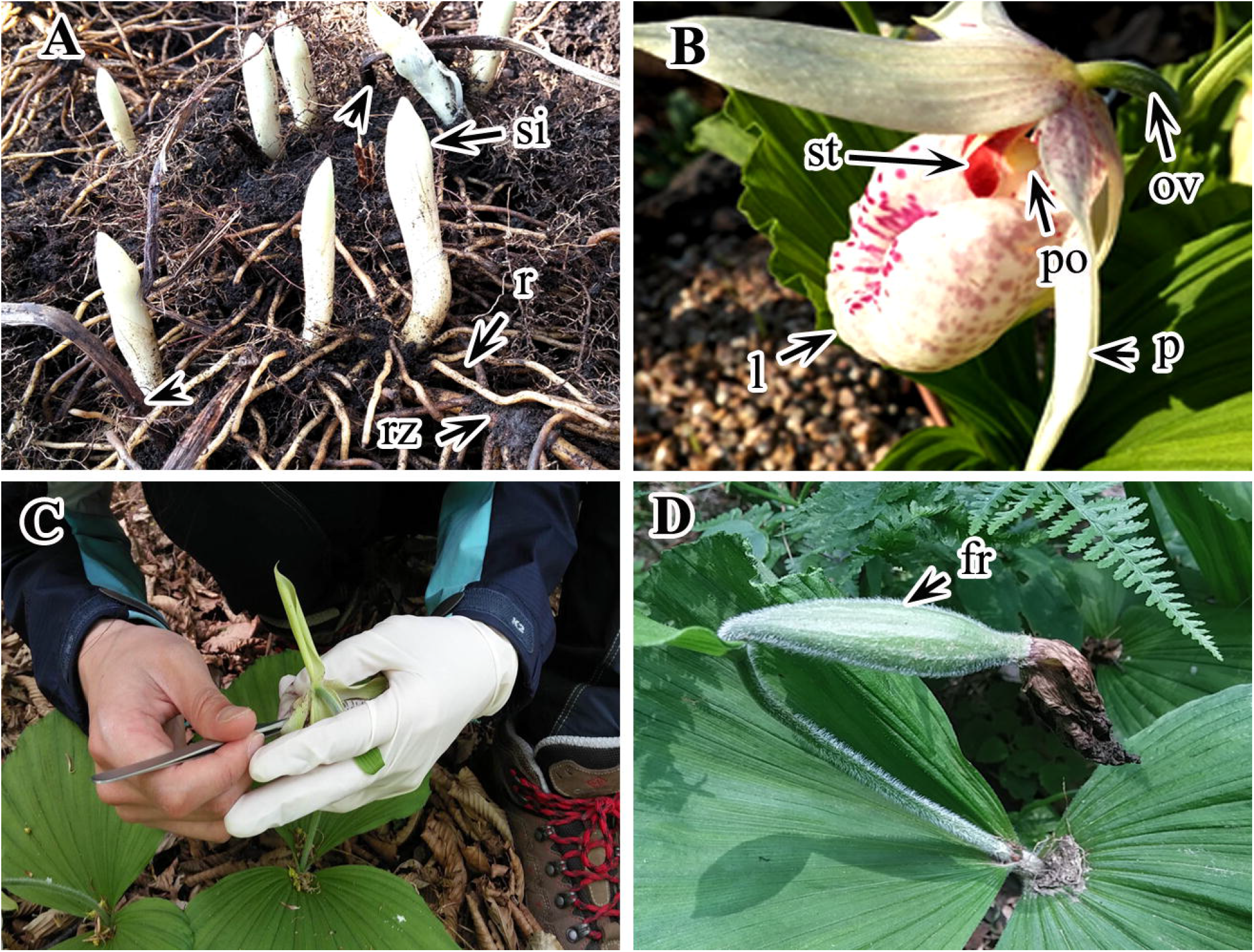
Vegetative and reproductive structure of *Cypripedium japonicum*. A. Young plant buds with rhizome and rhizoids. B. Open flower. C. Manual pollination of flower. D. Fruit. *Abbreviations*: fr, fruit; l, labellum; ov, ovary; p, petal; po, pollinia; r, rhizoids; rz, rhizome; si, shoot initial; st, staminoid.

### Microsporogenesis and the male gametophyte

The anther is bilobed and tetrasporangiate (Fig. 2A, 3A). Prior to maturation, the anther wall comprises five or six layers: an epidermis, an endothecium, two or three middle layers, and a tapetum (Fig. 2B). Unfortunately, due to a lack of young anthers we could not confirm the process of anther wall development. The tapetum is glandular and its cells are uninucleate throughout the development process (Figs. 2B, D). The middle layers degenerate at the same time as the spore tetrads give rise to free spores. Meanwhile, radial expansion is followed by the formation of wall fibers in the endothecium cells (Fig. 2E, H). The nuclei of the endothecium cells soon degenerate and individual cells become highly vacuolated. As the anther gets expanded the epidermal cells elongate tangentially and their nuclei also degenerate, before or after spore formation. Both epidermis and endothecium are persistent (Figs. 2G, H). In mature anther the epidermis is papillate and becomes sticky during anthesis whereas the endothecium develops fibrous thickenings (Fig. 2H).

**Fig. 2.**
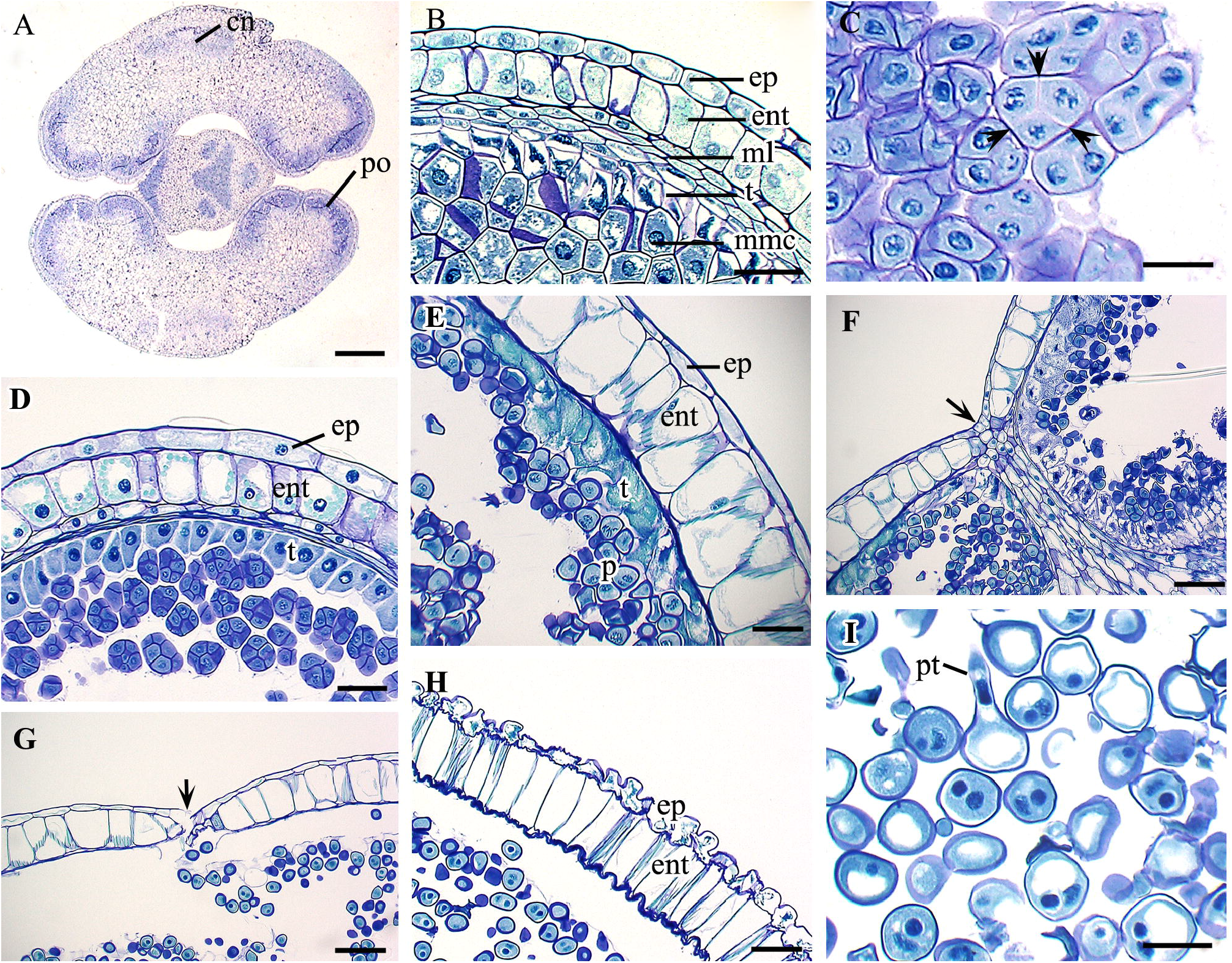
Development of anther and pollen grain in *Cypripedium japonicum*. A. Cross section (CS) of young pollinia. B. CS of young pollinia showing wall layers. C. Cytokinesis in microspores. D. CS of pollinia showing anther wall and microspore tetrads. E. CS of mature pollinia with fibrous endothecium and pollen grains. F, G. CS of mature pollinia showing dehiscing junction. H. Mature anther wall with papillate epidermis and fibrous endothecium. I. Two nucleate mature pollen grains just before pollination. *Abbreviations*: ent, endothecium; ep, epidermis; ml, middle layer; mmc, microspore mother cell; po, pollinia; pt, pollen tube; t, tapetum. Scale bars: A=500 µm, B, D=50 µm, C, I=30 µm, E, F, G, H=100 µm.

**Fig. 3.**
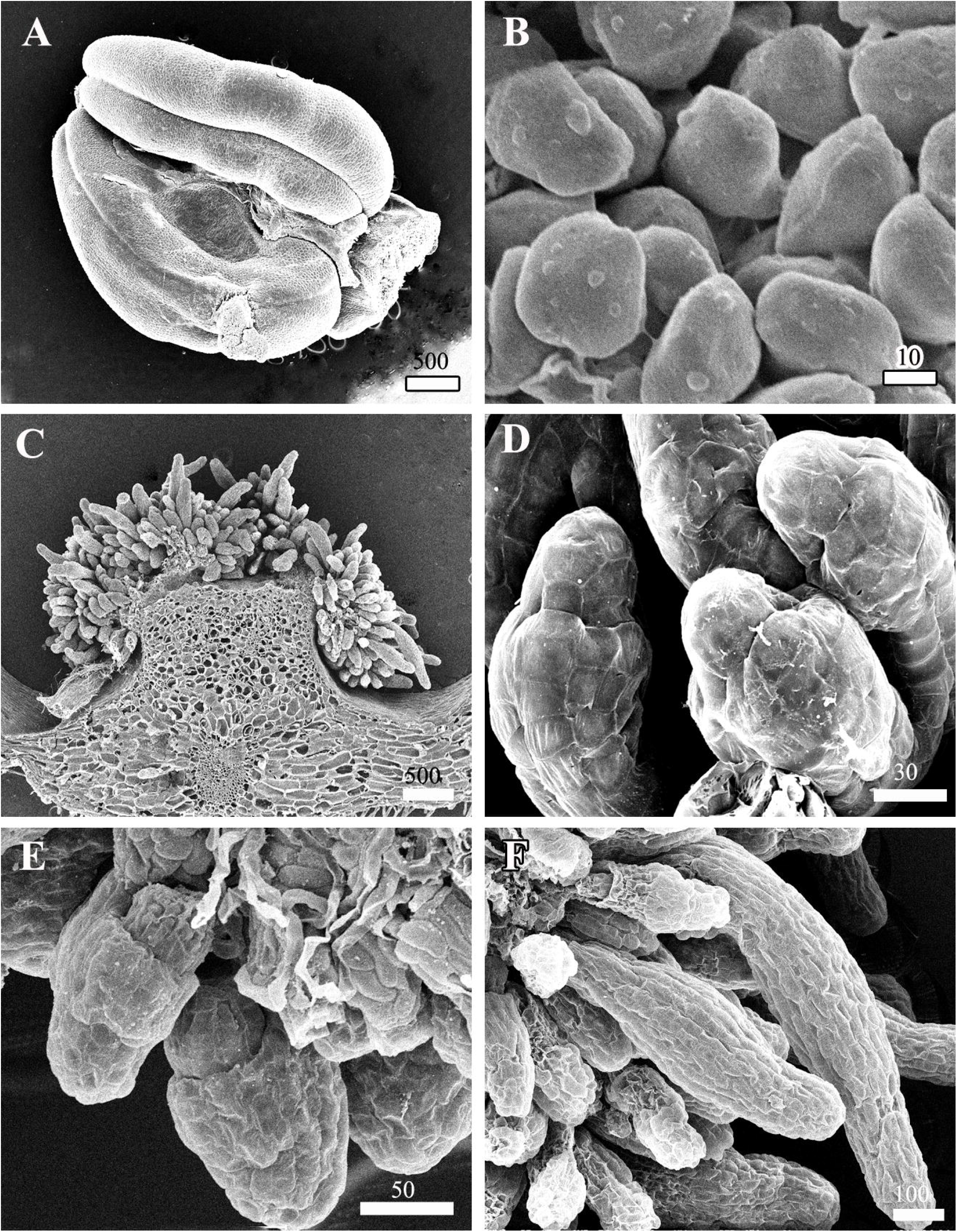
Scanning electron micrograph of pollinia and ovule of *Cypripedium japonicum*. A. Pollinia. B. Mature pollen grains. C. CS of ovule through placental ridge. D-E. Young ovular filament showing initiation of integuments and pollen tube. F. Young seed.

The sporogenus cells form an arc-like structure in the anther lobe and differentiate into microspore mother cells (Fig. 2B). The microspore mother cells undergo meiotic division, resulting in microspores. Cytokinesis in the microspore tetrad is always simultaneous and the resultant tetrad is tetrahedral, isobilateral, T-shaped, or linear (Fig. 2C, D). Unequal mitosis division occurs in the microspore, resulting in a larger generative cell and a smaller vegetative cell; thus pollen grains are binucleate during anthesis (Figs. 2H, I). No further division occurs before pollination although some pollen grains develop a pollen tube inside the anther (Fig. 2I). Pollen grains are single, non-aperturate, and smooth-walled (Fig. 3B). Anther dehiscence takes place from a common longitudinal slit between two chambers of the same lobe (Figs. 2F, G).

### Ovary

Flowers are inferior and ovaries are long, slender, and unilocular (Figs. 1B, 4A). In cross-section the ovary is circular in outline. Its wall is very thick and numerous multicellular hairs develop from the outer epidermis of the ovary (Fig. 4A). There are at least six large vascular bundles running through the massive parenchymatous cortical region and supporting the ovary of *C. japonicum* (Figs. 3C, 4A). The innermost layer of the ovule is meristematic in nature and acts as a placental region. During anthesis, the placental tissue within the inner wall of the ovary comprises three forked ridges representing three fused carpels (Fig. 3C). Each of these ridges had numerous profusely branched and uniseriate nucellar filaments (Fig. 3C-F).

**Fig. 4.**
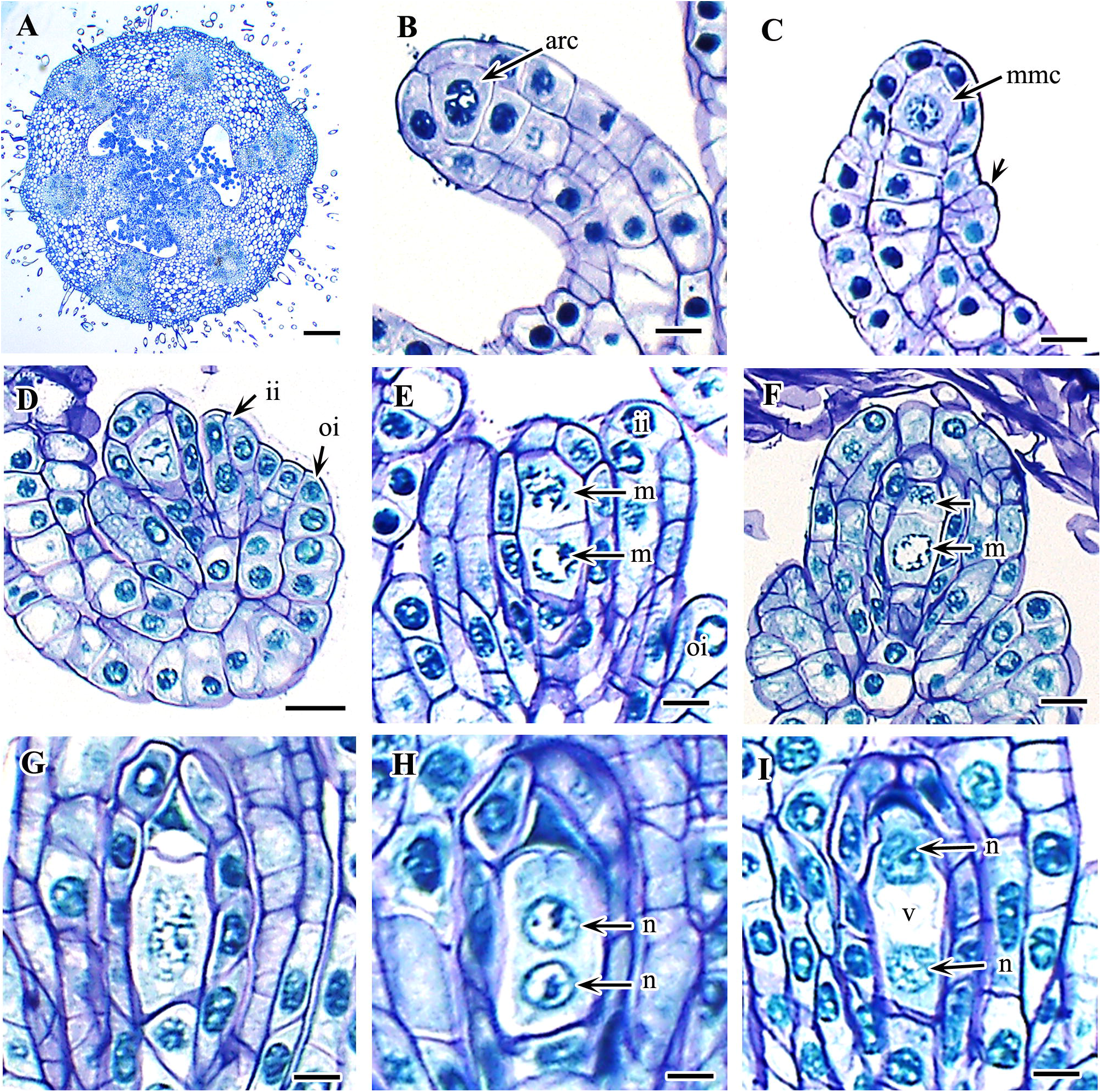
Development of ovule and megagametophyte in *Cypripedium japonicum*. A. Cross section of young ovary. B. Young ovular filament with archesporial cell. B. Young ovule with megaspore mother cell (MMC) and initiation of integument. D. Anatropous, bitegmic ovule. E. Mitotic division in MMC. F. Two unequal megaspore. G. Elongated functional megaspore. H–I. Two nucleate megaspore. *Abbreviation*: ii, inner integument; m, megaspore; mmc, megaspore mother cells; n, nucleus; oi, outer integument; v, vacuole. Scale bars: A=500 µm, B, C, E–I=20 µm, D=50 µm.

### Ovule

Each of the branched nucellar filaments which are raised in the placental ridge of the ovary comprises approximately four to seven nucellar cells surrounded by a single layer of nucellar epidermis (Fig. 4B). The topmost nucellar cell enlarges in size and functions as an archesporial cell. Meanwhile, integuments start to initiate in the nucellar filament. The origin of integuments in the ovule premordium is dermal (Figs. 3D, F; 4C). The inner integument differentiates first, becomes two-cell-layered, and eventually forms the micropyle before fertilization. The outer integument differentiates a little later but in a similar fashion to the inner integument, and grows beyond the inner integument after fertilization. During this period nucellar filaments regularly proliferate and begin to bend back towards the placental ridge and finally undertake an anatropous condition. The mature ovule is therefore tenninucellate, bitegmic, and anatropous in nature.

### Megasporogenesis and female gametophyte

The topmost cell of the nucellar filament differentiates into an archesporial cell (Fig. 4B). The single archesporial cell increases in size and directly functions as an MMC (Fig. 4C). Further enlargement of the MMC is followed by mitotic division to form two dyad cells of which only the chalazal megaspore is functional (Fig. 4D-F). The micropylar megaspore soon degenerates and the entire embryo sac development takes place from the chalazal megaspore. Together with degeneration of the micropylar megaspore the functional megaspore enlarges in size and becomes elongated (Fig. 4G). The first mitotic division in the chalazal megaspore produces a primary micropylar nucleus and a primary chalazal nucleus. The first division is immediately followed by the shifting of the nuclei to opposite poles of the cell (Fig. 4H, I). This shift in the primary micropylar and chalazal nuclei creates a large central vacuole in the premature embryo sac (Figs. 4I, 5A). The second mitotic division occurs in the primary micropylar nucleus and primary chalazal nucleus to form a four-nucleate embryo sac with two nuclei at each pole (Fig. 5B). At this stage, two micropylar nuclei divide once again to form four nuclei, but no further division is observed in the two chalazal nuclei (Fig. 5C). Thus the mature embryo sac contains six nuclei (Fig 5D). The four micropylar nuclei form an egg, two synergids, and one polar nucleus whereas the two chalazal nuclei form the polar nucleus and an antipodal (Fig. 5E). The antipodal cells are ephemeral and degenerate before or after fertilization.

**Fig. 5.**
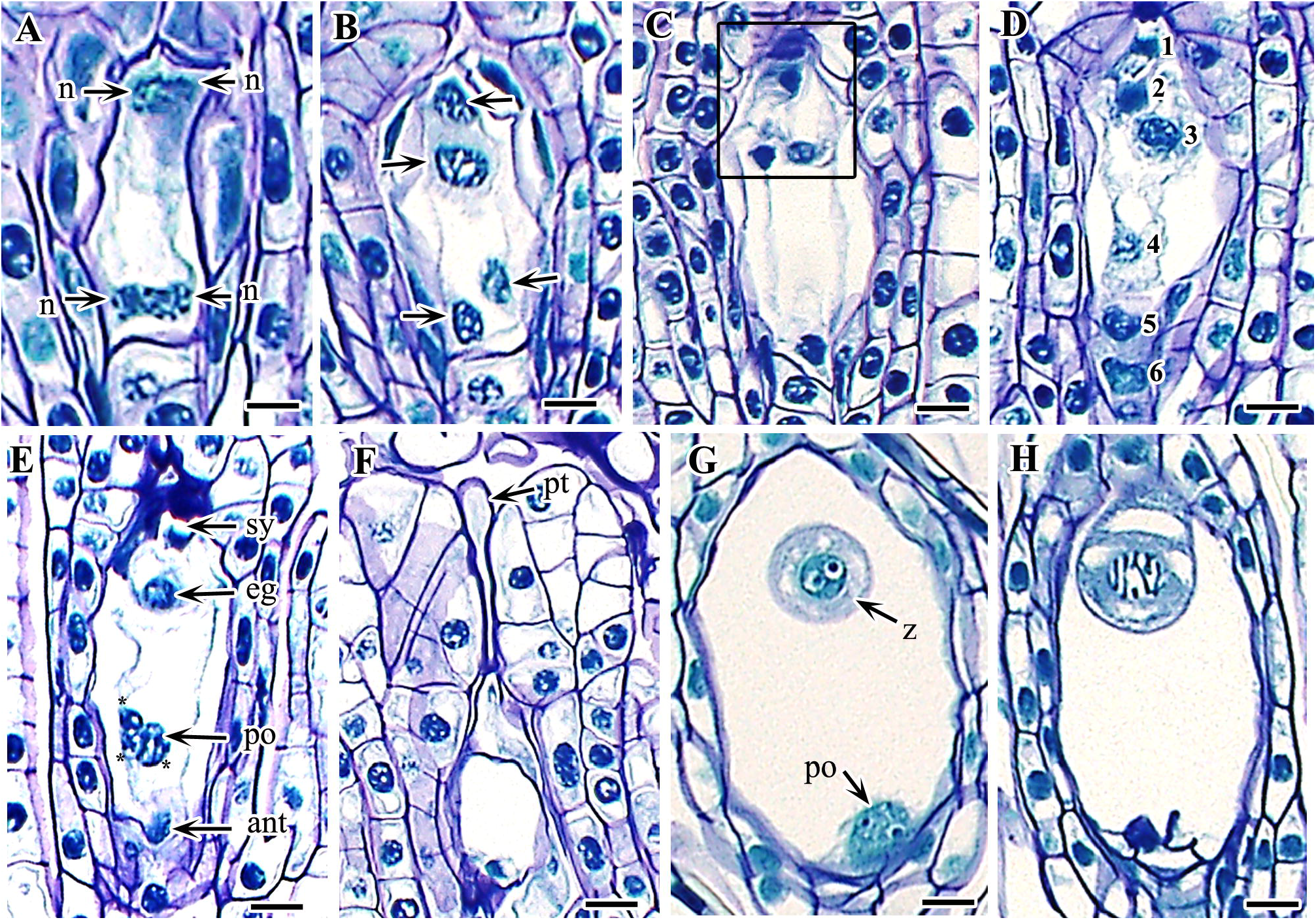
Development megagametophyte and zygote in *Cypripedium japonicum*. A. Two nucleate embryo sac, nuclei undergoing first mitotic division. B. Four nucleate embryo sac. C. Micropylar nuclei undergoing second mitotic division (rectangle line). D–E. Six nucleate mature embryo sac (asterisks in central cell of E indicate triple fusion). F. Pollen tube. G. Zygote and endosperm nucleus. H. Proembryo. *Abbreviations:* ant, antipodal cell; eg, egg; n, megaspore nucleus; po, polar nuclei; pt, pollen tube; sy, synergid; z, zygote. Scale bars: A–H=20 µm

### Pollination and fertilization

The anther matures much earlier than the ovule. The ripened anther is yellow with a shiny, sticky surface. The flowers were hand-pollinated during May 2–5. The sticky surface of the anther helps to attach it on the stigma. During pollination pollen grains were binucleate and the pollen tube was about to grow. In contrast, the nucellar filaments were just ready or still not ready to organize the archesporial cells. Fertilization occurs almost 7–8 weeks after pollination. An entangled mass of pollen tubes was observed across the placental ridges when the embryo sac was in two nuclear stages. Fertilization is porogamous: the pollen tube enters the embryo sac through the micropyle, resulting in a diploid embryonic zygote by fusion of one male gamete with an egg, and a triploid endosperm zygote after the fusion of one male gamete with the diploid polar nucleus (Fig. 3E, 5F, G). However, the endosperm zygote does not develop further and degenerates before the embryo occupies the whole space.

### Embryogenesis

The zygote, the precursor of embryo formation, is globular (Fig 5G, H). The first division of the zygote is always transverse, resulting equal-sized basal and terminal cells (Fig. 6A). Mitotic division is immediately followed by rapid elongation of the basal cells, which acquire a large vacuole; in contrast, terminal cells remain unchanged, with a dense cytoplasm (Fig. 6B). Both cells divide transversely producing a four-celled linear pro-embryo (Fig. 6C, D). No further division was observed in the two basal cells, which directly function as the suspensor whereas the two terminal cells, after a series of divisions, produce the whole embryonic body. Out of the two terminal cells, transverse division occurs in the sub-terminal cell and longitudinal division occurs in the terminal one, and these four, eventually gave rise to the embryo proper (Fig. 6 E, F). Each of the terminal cells is then divided by a vertical wall at right angles to the first one, and produces a quadrate, whereas the upper tire of the sub-terminal cell divides by longitudinal division (Fig. 6G). The lower tire cell of the embryo proper soon undergoes longitudinal division resulting in a 10-celled embryo proper, including the two suspensor cells (Fig. 6H). During the early globular stage, additional cell divisions occur in the terminal tire and upper sub-terminal tire resulting in a 16-celled embryo proper (Fig. 6I). Periclinal divisions from the outer layer of cells give rise to protoderm (Fig. 6J, K, L). The suspensor was inconspicuous and represented by two cells. Normal mitotic division occurs in the suspensor, initially forming two cells which are without any special modification or development, and which directly function as a suspensor. Cleavage polyembryony was also observed, although its frequency was quite low (Fig. 7A-C). Polyembryony is due to the proliferation of the zygote and both embryos developed in a similar fashion to the normal embryo.

**Fig. 6.**
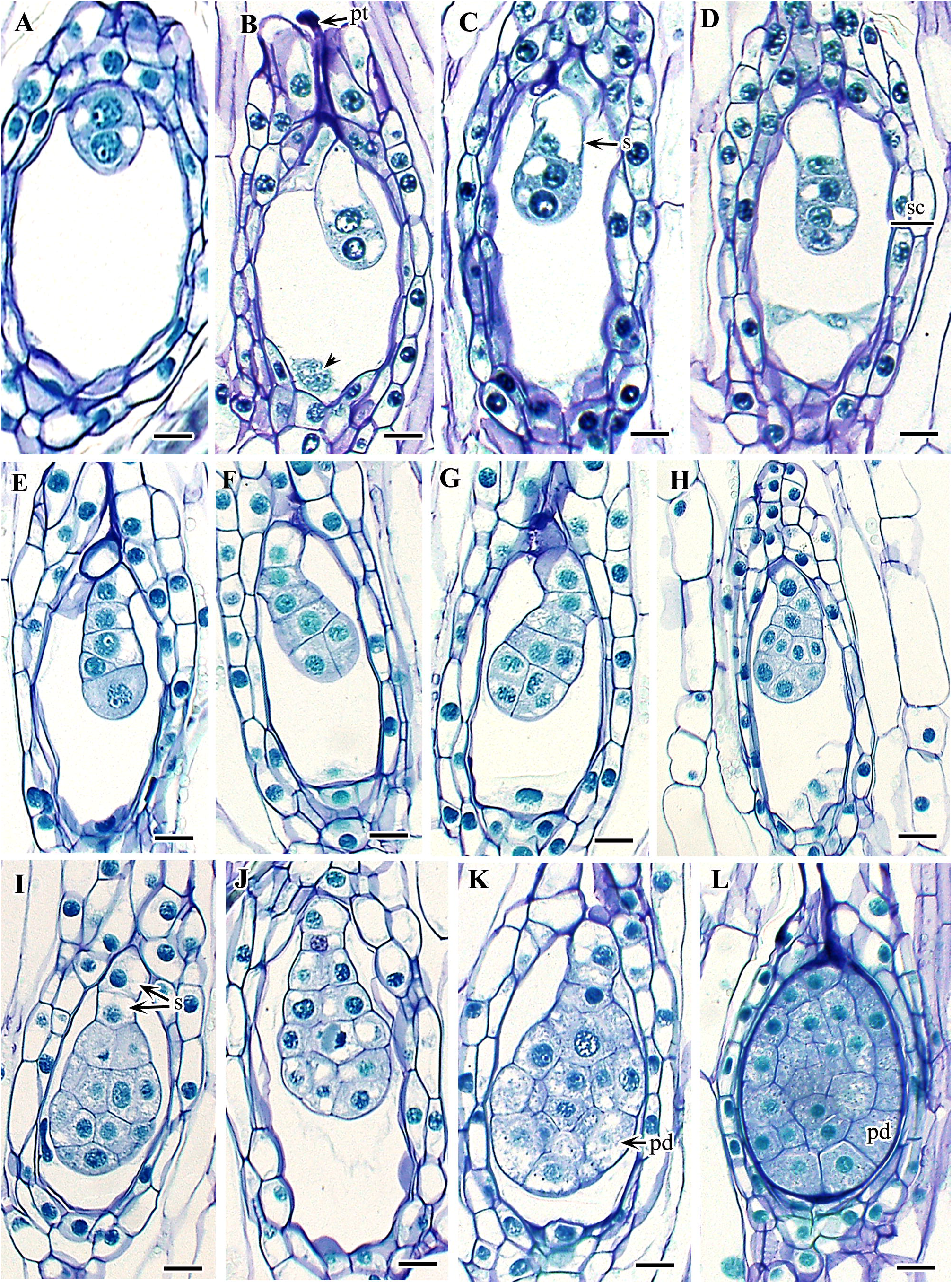
Development of embryo in *Cypripedium japonicum*. A. First transverse division in zygote forming two, terminal and basal, celled pro-embryo. B. Elongation of basal cell forming unequal size of two embryonal cells. C. Second transverse division in terminal cell forming three celled pro-embryo (terminal, sub-terminal, and basal). D. Third division in terminal cell forming four-celled linear pro-embryo. E. Five celled linear pro-embryo (basal cell divides in this stage forming two celled suspensor). F. Longitudinal division in terminal cell of five celled embryo showing two juxtaposed cells. G–I. Pro-embryo forming early stage of globular pro-embryo (endosperm nucleus completely degenerate after this stage). J–H. Periclinal division in embryonic cell (except two basal suspensor cells) forming protoderm layer of embryo. L. Fully developed embryo covering whole inner space of the seed. *Abbreviations*: pd, protoderm layer; pt, pollen tube; s, suspensor; sc, seed coat. Scale bars: A–L=20 µm

**Fig. 7.**
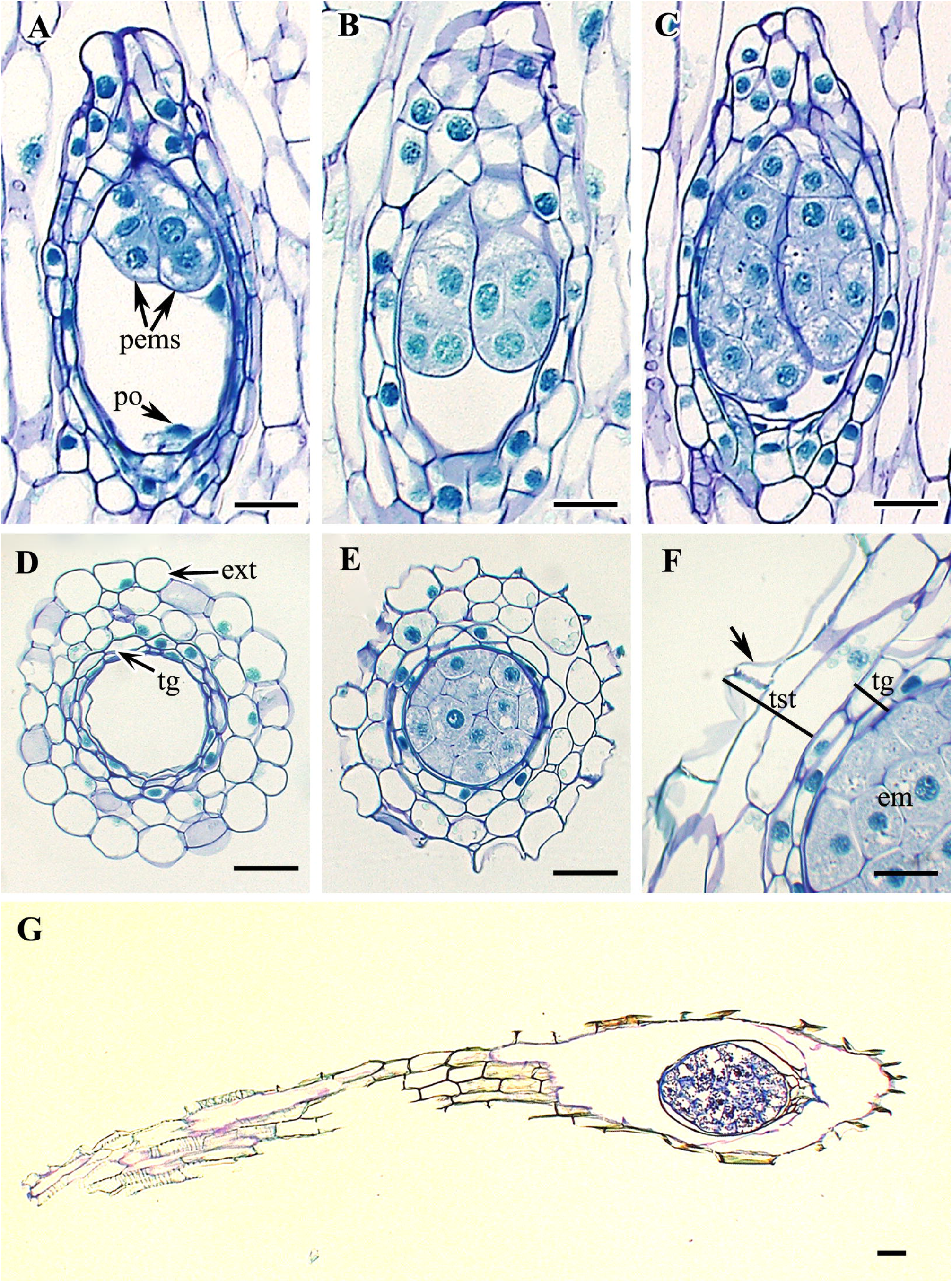
Polyembryony and seed coat development in *Cypripedium japonicum*. A–C. Cleavage polyembryony in different stages. D-F. Transverse division of seed showing seed coat development. G. Longitudinal section of mature seed. *Abbreviations*: em, embryo; ext, exotesta; pems, proembryos; po, polaaar nuclei; tg, tegmen; tst, testa. Scale bars: A–E=20 µm, F=10 µm, G=30 µm.

### Fruit

The capsule of *C. japonicum* is elongated, and fusiform to spindle-shaped in shape (Fig 3D). The size range is 5±0.5 cm in length by 1.5±0.3 in diameter. The surface is covered by numerous, multicellular, transparent hairs. The remaining petals can be seen at the tip of the mature fruit. Seeds are dispersed from a longitudinal slit in the dry and ripened fruit.

### Seeds

The mature seeds are brown, minute, narrowly elliptical, and curved. Size range is 2.5±0.3 mm in length by 0.15±0.07 mm in width. Prior to maturity, the seed coat is four- or five-cell-layered. The testa comprises two or three cell layers and tegmen is composed of two layers. The outermost exotestal cells are comparatively large, and rounded to oval or irregular-shaped in cross-section. The exotesta is followed by a meso- and then an endotestal layer. Both these layers are composed of similar types of cells to the exotesta but are smaller in size. The innermost two layers represent the tegmen, cells of which are smaller and elongated in cross-section (Fig. 7D–G). During maturity the exoetsta becomes papillate and the remaining layers degenerate and entire seed coat is represented by thin, papery layers.

### Reproductive calendar

The reproductive cycle of *C. japonicum* begins under the soil. All the developmental stages with time frame have been provided in Table 1 and Fig. 8. Pollinia development starts during mid-March when the whole plant lies below the soil surface. The primary sporogenous cells directly function as pollen mother cells and this process occurs from late March until early April. Meiosis in the pollen mother cells results in variously shaped pollen tetrads by the middle of the April. The nuclei of pollen grains divide by mitotic division during late April and pollen grains are ready for pollination by early May. In our study population the optimum period for pollination was May 2–5. Ovular development starts about 2 weeks before pollination. The archesporial cell appears during the third week of April and lasts until about the middle of May. The MMC was observed at 5–7 days after pollination and lasted until the last week of May. Meiosis in the MMC starts 2 weeks after pollination and degeneration of the micropylar cell, followed shortly by first nuclear division, occurs almost 4 weeks after pollination. The second and third nuclear divisions occur 36–45 days after pollination. Thus in *C. japonicum* the female gametophyte matures 5–7 weeks after pollination. Fertilization was observed during late June to early July, about 42–48 days after pollination, whereas the zygote started to divide by early July and a two-to-four-celled linear pro-embryo was formed about 55–60 days after pollination. The constant cell division in the four-celled pro-embryo resulted in an eight- and then 16-celled globular embryo about 65 days after pollination. Within 72–80 days whole interior of the seed was filled with embryo. Mature seeds were harvested 90 days after pollination.

**Fig. 8.**
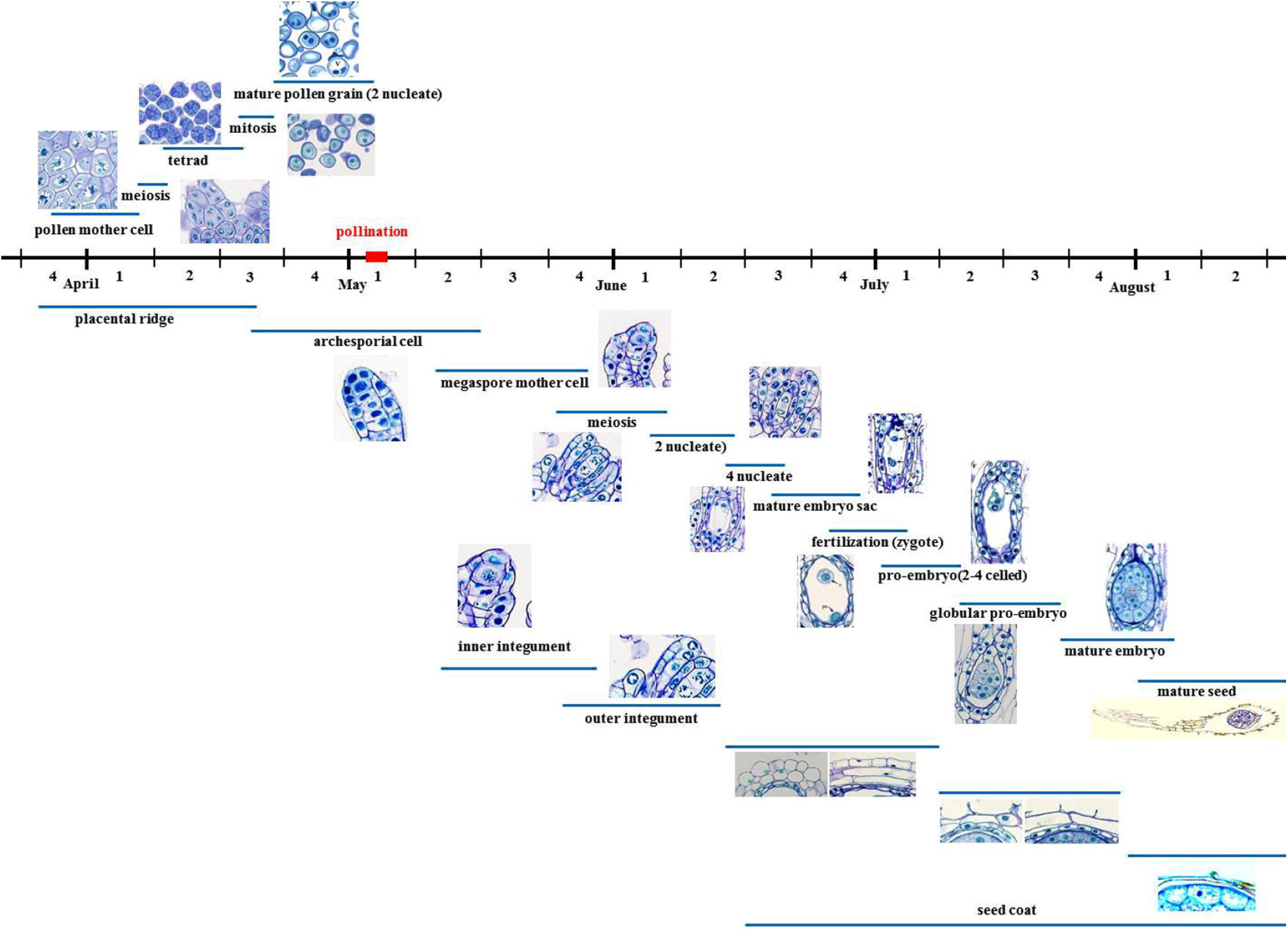
Reproductive calendar of *Cypripedium japonicum*. Blue line indicating the duration for the corresponding stage. Numbers indicating weeks of the corresponding month.

## Discussion

This study provides a complete overview of the reproductive structures and embryological organization of the endangered slipper orchid, *C. japonicum*. In nature, this non-rewarding orchid reproduces by both sexual and vegetative methods. However, the general trend of fruit set for non-rewarding orchids including *C. japonicum* in natural conditions is very low (Tremblay *et al*., 2005; Suetsugu and Fukishima, 2013). It is quite understandable that such non-rewarding species is often challenged by pollinator limitation and thus exhibits low reproductive success in nature (Zimmerman and Aide, 1989; Calvo and Horvitz, 1990; Neiland and Wilcock, 1998; Tremblay *et al*., 2005; Sun *et al*., 2009). Suetsugu and Fukishima (2013) found the fruit set of *C. japonicum* by natural pollination was only 14.9%, whereas it was even worse (i.e. only 5.2–7.7%) in the samples of Sun *et al*. (2009). Although we carried out hand pollination in our study population, we observed very low fruit set (≤5%), almost congruent with Sun *et al*. (2009), in naturally pollinated populations near our sampling site. The fruit set in hand-pollinated populations in our experimental site was much higher (60–65%), more than 12-fold local the naturally pollinated population. Hand-pollination experiments have already confirmed self-compatibility in *C. japonicum* (Sun *et al*., 2009; Suetsugu and Fukishima, 2013); however, we performed cross-pollination between individuals of both the same and different clumps of the population.

A majority of recent embryological studies of the family Orchidaceae have focused on ovule, embryo, and suspensor development. Pollinia and male gametophyte developmental studies are scarce for orchids and only a few studies are available on *Spiranthes* and *Ophrys* species (Akbeke, 2012; Kant *et al*., 2013; Wang *et al*. 2016). Before that, Sood and Rao (1988) studied embryology of *C. cordigerum*. This study attempted to provide a comprehensive statement on embryology including pollinia structure and male gametophyte development in *C. japonicum*. The flowers of *C. japonicum* are protandrus: the male gamete develops and matures much earlier than the ovule. Protandry in orchids was first observed in *Spiranthes* and *Listea* (Darwin, 1862) and is now spread across the tribes Cranichideae, Orchideae, Neottieae, and Cymbidieae (Darwin, 1862; Ackerman, 1975, 1977; Catling, 1983; Sipes and Tepedino, 1995; Singer and Sazima, 2001a, 2001b; Singer, 2002; Singer and Koehler, 2003). In *C. japonicum* anther development is underway (possibly during mid March) even when the entire plant remains underneath the frozen soil surface. In previous studies, the monocotyledon type of anther wall formation has been observed in some orchid species (Prakash and Lee, 1973; Sood and Rao, 1986; Sharma and Vij, 1987); however, due to a lack of pollinia in the early developmental stages, wall development in *C. japonicum* could not be recognized here. Nevertheless, the anther wall, from the beginning of microspore mother cell development to the end of microsprogensis, was six- or seven-cell layered. This is in agreement with Sood and Rao (1988), who mentioned six to eight layers in *C. cordigerum*, although Wirth and Withner (1959) reported a five-cell-layered anther wall in most of the terrestrial orchids. In contrast, Aybeke (2012) found only four-layered anther walls in *Ophrys mammosa*. In this study, we found a single-layered tapetum except in places where a few cells divide periclinally, making two layers. A partly two-layered tapetum in the area between the two microsporangia has been described for *Ophrys mammosa* (Aybeke, 2012). In contrast, Sood and Rao (1988) described a two- or three-layered tapetum in *C. cordigerum* which is infrequent for most of the orchids (Johri *et al*., 1992). More interestingly, Sood and Rao (1988) revealed two-layered fibrous thickening in the mature anther wall of *C. cordigerum*, formed by an edothecial and the outermost middle layer. However, in *C. japonicum* only the endothecial layer develops such thickenings and none of the remaining middle layers persist in the mature anther. Moreover, none of the previous works have mentioned double-layered wall thickening in the family (Wirth and Withner, 1959; Sharma and Vij, 1987; Rao, 1967; Karanth *et al*., 1979; Sood and Rao, 1986; Vij and Sharma, 1987; Johri *et al*., 1992; Aybeke, 2012). This disparity in nature and number of wall layers makes this feature a worthwhile taxonomic and embryological development criterion in Orchidaceae (Swamy, 1949a; Sood and Sham, 1987; Aybeke, 2012).

Early studies suggested that a simultaneous type of cytokinesis in microspores is the characteristic feature of the family Orchadaceae (Swamy, 1949a; Johri *et al*., 1992); however, successive types are also often observed in this family (Aybeke, 2012; Wang *et al*., 2016). In this study, we found a simultaneous type of microsporogenesis in *C. japonica*. This is in agreement with Sood and Rao (1988) and Poddubnaya-Arnold (1967) who described simultaneous microspore development in other *Cypripedium* species, although before that Guignard (1882) reported the successive type for this genus. This variation of cytokinesis in *Cypripedium* species might not be surprising as both types of cytokinesis have been observed in *Spiranthes sinensis* (Kant *et al*., 2013; Wang *et al*., 2016), and also in a few other angiosperms, for instance *Rauvolfia canescene* (Mayer, 1938), *R. serpentina* (Ghimire *et al*., 2011), and *Catharanthus pusilis* (Bhasin, 1971). We observed that the resultant pollen tetrads are isobilateral, tetrahedral, linear, or T-shaped, which are common tetrad types in the family, although only decussate, isobilateral, and tetrahedral types has been observed in *C. cordigerum* (Sood and Rao, 1988). The mature pollen grains are exclusively single, non-aperturate, and two-celled in *C. japonicum* as was found in other *Cypripedium* spp. (Poddubnaya-Arnoldi, 1967; Sood and Rao, 1988). The ovule, embryo sac, and megagametophyte development in *C. japonicum* follows the usual pattern as described in other Orchidaceae. The important embryological processes we observed in this study included movement of nuclei towards the micropylar and chalazal end during the binucleate stage and formation of a large central vacuole; re-fusal in the second nuclear divisions by the chalazal nuclei, resulting in a six-nucleate mature embryo sac; single-celled ephemeral antipodal; triple fusion and a binucleate endosperm cell; a one- or two-celled suspensor; and cleavage polyembryony.

The development of integument starts when the archesporial cell enlarges and forms the MMC. During this time a couple of dermal cells of the ovular filament just below the MMC increase their size and divide by a transverse wall. Thus both the integuments are dermal in origin as described in *C. cordigerum* (Sood and Rao, 1988) and inner integument is usually involved in micropyle formation. Strangely, Poddubnaya-Arnoldi (1967) reported a unitegmic ovule in *C. insigne* which is not common in the genus, even in the family Orchidaceae (Davis, 1966; Johri *et al*., 1992). Initially, the division of MMC usually results in an equal-sized dyad; however, in this species only the chalazal one proliferates and becomes functional, which results in development of the whole embryo sac, while the micropylar cell soon degenerates. Pace (1907), who observed fertilization and embryogenesis in four *Cypriperdium* species, described the four-nucleate bisporic embryo sac as characteristic of the genus *Cypripedium* and thus termed it the ‘*Cypripedium* type’ of embryo sac. However, this was later corrected by other researchers who found five- to eight-nucleate bisporic embryo sacs instead of four-nucleate ones in different *Cypripedium* species (Prosina, 1930; Carlson, 1940, 1945; Swamy, 1945; Poddubnaya-Arnoldi; 1967; Kimura, 1967; Sood and Rao, 1988). The results of this study are relatively close to the latter idea, observing as we did a six-nucleate bisporic mature embryo sac. After the four-nucleate embryo sac is formed, only two micropylar nuclei divide futher and form four nuclei but the chalazal nuclei fail to divide and thus the mature embryo sac comprises six nuclei. In this case the antipodal is represented by a single cell because one nucleus from each side moves to the center of the embryo sac and forms a binucleate polar cell which produces the endosperm cell after triple fusion. This failure of subsequent nuclear division in chalazal nuclei is termed the ‘strike’ phenomenon. This rare feature has been observed in *Paphiopedilum* spp. in which one of the micropylar and two chalazal nuclei of the four-nucleate-stage embryo sac fail to divide and thus mature embryo sac comprises five nuclei (Francini, 1931).

The pollen tube enters through the micropyle and fertilization likely occurs between 7 and 8 weeks after pollination. We confirm that double fertilization and triple fusion is evidently occurring in *C. japonicum*, although the primary endosperm is binucleate and short-lived. The degeneration of endosperm nuclei occurs before or after the embryo has reached the eight-celled stage. Previously, Pace (1907) explained the common occurrence of two (or, rarely, four) endosperm nuclei in *Cypripedium* spp., but no triple fusion was properly observed. According to Pace (1907), the primary endosperm is formed by the fusion of the polar nucleus, one synergid, and one male nucleus but this study rejects this fact for *C. japonicum*. Later, Sood and Rao (1988) also found a binucleate primary endosperm in *C. cordegerum* whereas Poddubnaya-Arnoldi (1967) recorded a six-nucleate endosperm for *C. insigne*. The maximum number of endosperm nuclei recorded in the family is 12 in *Vanilla* and 16 in *Galeola septentrionalis* (Chugai, 1971). Moreover, some previous studies disputed the double fertilization in orchid. For instance, according to Savina (1978) in *Listera ovata* and *Ophrys insectifera*, only one male gamete fuses with the egg but a second male gamete does not fuse with the secondary nucleus, thus no triple fusion occurs. Correspondingly, Maheshwari and Narayanaswami (1952) reported only one sperm nucleus being released from the pollen tube in *Spiranthes australis*; thus no fertilization of polar nuclei is likely to happen in that species. Tarasaka *et al*. (1979) suggested the failure of generative cell division in pollen, resulting in only one sperm cell in *S. sinensis* and thus no male gamete left for triple fusion; however, this was later denied by Battaglia (1980). Recently, *in vitro* pollen germination of *S. sinensis* by Wang *et al*. (2016) found two male gametes in the pollen tube and suggested that triple fusion in this species is expected.

One of the most notable features during the course of embryo development in the *Cypripedium* is lack of an elaborated suspensor cell. The suspensor is likely to be one- or two-celled in *C. japonicum*. Previous studies suggested that the suspensor in *Cypripedium* is either totally absent (Treube, 1879; Schnarf, 1931) or represented by an inconspicuous one or two cells (Treub, 1879; Pace, 1907; Carlson, 1940; Sood and Rao, 1988). It is interesting that majority of structural studies of embryo development in Orchidaceae have focused on suspensor development; even Swamy’s (1949b) embryo classification idea was based on the suspensor structure. According to Swamy’s five embryo categories based on suspensor morphology in Orchidaceae, *Cypripedium* belongs to the type I category, in which the suspensor initial cell does not undergo any division but remains without much elongation. Similar to *Cypripedium* species, Prakash and Lee (1973) and Tohda (1971) reported unicellular and vestigial suspensors in *Spathoglottis pliccata* and *Lecanorchis* spp. respectively. Moreover, unicellular suspensor in *Hetaeria shikokima* develops into a long projection through the micropyle, becomes twisted, and enters the tissue of the placenta (Tohda, 1967). The function of unique vacuolated suspensor of the orchid remains uncertain, although Lee *et al*. (2006) found specific structural specialization with possible ‘transfer cell’ morphology in *Paphiopedilum delentii*. However, this is yet unclear for the taxa like *Cypripedium* in which the specialized suspensor is lacking. Thus, in order to understand the definite structural specialization of suspensor in such species further studies focusing on ultrastructure of suspensor is required.

The first division of the zygote in *C. japonicum* follows a regular pattern, forming a terminal cell and a basal cell. Swamy (1949b) classified orchid embryos into three types based on the embryonic cell division pattern. According to the Swmy’s report *Cypripedium* belongs to group A, based on the cell division pattern of the developing embryo and type I based on the suspensor morphology. However, Sood and Rao (1988) categorized the embryogeny of *C. cordigerum* under group B of Swamy’s (1949b) classification, stating a greater share of basal cell in the organization of the mature embryo. Before Swamy’s classification, Carlson (1940) in *C. parviflorum* reported a one-celled suspensor originating from the basal cell and the entire embryo developing from the terminal cell alone. The result of this study fundamentally agreed with Carlson (1940) rather than Swamy (1949b) and Sood and Rao (1988) as the basal cell do not takes part in the formation of embryonal mass in *C. japonicum*. One of the most notable findings in *C. japonicum* is that the second and third divisions in the terminal cell also occur by transverse wall formation, resulting in a five-celled linear pro-embryo. The longitudinal division occurs in the terminal cell of the five-celled pro-embryo, forming two juxtaposed cells. Meanwhile, the basal cell divides transversely, forming a two-celled suspensor. From the results of this study we confirm that *C. japonicum* does not follow the pattern of embryo and suspensor development described earlier in this genus.

One of the major threats to decreasing populations of *C. japonicum* is the complex pollination mechanism and low fruit set under natural conditions. Although *in vitro* seed germination has received the most attention for different *Cypripedium* species, no successuful *in vitro* germination methods have been suggested for *C. japonicum*. Results of germination studies for the genus *Cypripedium* have been contradictory and, in many cases, somewhat empirical in nature, with limited control of variability (Pauw and Remphrey, 1993). Several reports have suggested that immature seed improved the *in vitro* germination compared to mature seed in *Cypripedium* species (Witthner, 1953; Pauw and Remphrey, 1993, Jiang *et al*., 2017). The reproductive cycle presented in this study might be useful in future studies to select the appropriate embryonic stage for germination experiment. According to Pauw and Remphery (1993), seed collected after 8 weeks after pollination (WAP) sharply decreased the germination percentage in three *Cypripedium* species. Recently, Jiang *et al*. (2017) found the optimum germination percentage in *C. lentiginosum* to be from seeds collected at 90–105 days after pollination (DAP), at which time the embryo is in early globular to globular stage. In *C. japonicum* the embryo in the early globular to globular stage was observed at 9– 10 WAP or 65–75 DAP. Thus, the seeds collected near to this period could be useful for *in vitro* seed germination for this species. In conclusion, this study presents comprehensive embryological features of the endangered orchid *C. japonicum* and also provides a reproductive calendar for the species. Although, some previous investigations reported a number of embryological features for the genus *Cypripedium*, the data pertaining to *C. japonicum* are entirely new in this study. The findings of the present study agreed with the previously investigated species of the genus in several features but also contrast in some occasions, for instance, fibrous wall thickening in mature anther and tapetum structure with Sood and Rao (1988), and mature embryo sac, triple fusion, and embryo development with Pace (1907), Sood and Rao (1988), and Poddubnaya-Arnoldi (1967). The overall comparison of gametophyte and embryo development features suggested that the previous embryological reports on *Cypripedium* are not sufficient to characterize the entire genus. Moreover, some of the previously reported features are ambiguous and thus need to be corrected by new studies. This study helps to clarify some contradictory findings from previous reports. Given this, the results of this study will certainly be helpful for future research and will provide a fundamental reference point for the embryological data for the genus *Cypropedium* and likewise the family Orchidaceae. In addition, the reproductive cycle presented in the study will be helpful for plant breeders to understand the suitable embryo developmental stages for *in vitro* germination of *C. japonicum*.

## Acknowledgement

This study was financially supported by the project ‘Ex-situ Conservation of Forest Plant Seeds in Korea (KNA1-2-29).

